# Drug GRADE: an integrated analysis of population growth and cell death reveals drug-specific and cancer subtype-specific response profiles

**DOI:** 10.1101/2020.02.26.966689

**Authors:** Hannah R. Schwartz, Ryan Richards, Rachel E. Fontana, Anna J. Joyce, Megan E. Honeywell, Michael J. Lee

## Abstract

In the pre-clinical evaluation of anti-cancer drugs, two different measurement approaches are used: relative viability, which scores an amalgam of growth arrest and cell death, and fractional viability, which more specifically scores the degree of cell killing. In this study, we directly quantify relationships between drug-induced growth inhibition and drug-induced cell death by counting live and dead cells over time using quantitative microscopy. We find that most drugs affect both growth and death, but with different proportions and with different relative timing. These features lead to a non-uniform and unpredictable relationship between the canonical relative and fractional drug response measurements. To unify these disparate measurements, we create a new data visualization and data analysis platform, called drug GRADE, which characterizes the degree to which cell death contributes to an observed reduction in population size for any given drug. Our new method reveals both drug- and genotype-specific drug responses, which are not captured using traditional pharmaco-metrics. Taken together, this study highlights the extremely idiosyncratic nature of drug-induced growth and cell death and provides a new analysis tool for quantitatively evaluating these behaviors.

## INTRODUCTION

Precise evaluation of a cell’s response to drug is a critical step in pre-clinical drug development. Failures in this process have contributed to issues with irreproducibility of phenotypes across experimental platforms, spurious associations in precision medicine, and misannotated mechanisms of drug action (Bruno et al., 2017; Chopra et al., 2019; Hafner et al., 2019; Haibe-Kains et al., 2013). Recent studies continue to reveal that we generally do not know how drugs function, even for drugs that are well-studied and precisely engineered (Lin et al., 2019). Traditional methods to evaluate a drug response have relied on pharmacological measures of a drug’s dose-response relationship, such as the EC50 or IC50. These features are important, but they reveal a biased and incomplete insight. Notably, measures of drug potency such as the EC50/IC50 are poorly correlated with other important features, such as the maximum response to a drug (*i.e*. drug efficacy) (Fallahi-Sichani et al., 2013). Furthermore, measures of drug potency provide minimal insights into the mechanisms of drug action. In recent years several new drug scoring algorithms have been developed to improve the evaluation of pharmacological dose-responses. Methods have been developed that facilitate an integrated evaluation of drug potency and efficacy (Fallahi-Sichani et al., 2013; Meyer et al., 2019). Additionally, it has now been well-demonstrated that differences in growth rate between cell types were a confounding factor in most prior measurements of drug sensitivity (Hafner et al., 2016). Correcting for these artifactual differences in the apparent drug sensitivity generates a more rational evaluation of drug sensitivity and has identified drug sensitivity-geneotype relationships that are missed using traditional methods (Hafner et al., 2016; Harris et al., 2016).

One issue that has not been explored in detail is the underlying data itself. In nearly all cases, drug sensitivity is scored by comparing the relative number of live cells in the context of drug treatment to the number of live cells in a vehicle control condition. This metric is variably referred to as “relative viability”, “percent survival”, “percent viability”, “drug sensitivity”, “normalized cytotoxicity”, etc. (hereafter referred to as Relative Viability, or RV). RV is a convenient measure of drug response, and can be quantified using most commonly used population based assays (e.g. MTT, Cell Titer-Glo, Alomar blue, colony formation, etc.). Importantly, changes to relative viability can result from partial or complete growth arrest, increased cell death, or both of these behaviors (Hafner et al., 2016). Because relative viability is determined entirely from live cells, this measure provides no insights into the number of dead cells, or importantly, the relationship between growth arrest and cell death following application of a drug. When using RV, it is generally unclear to what extent a cell population is undergoing growth arrest versus cell death at a given drug concentration (Figure 1A).

**Figure 1:**
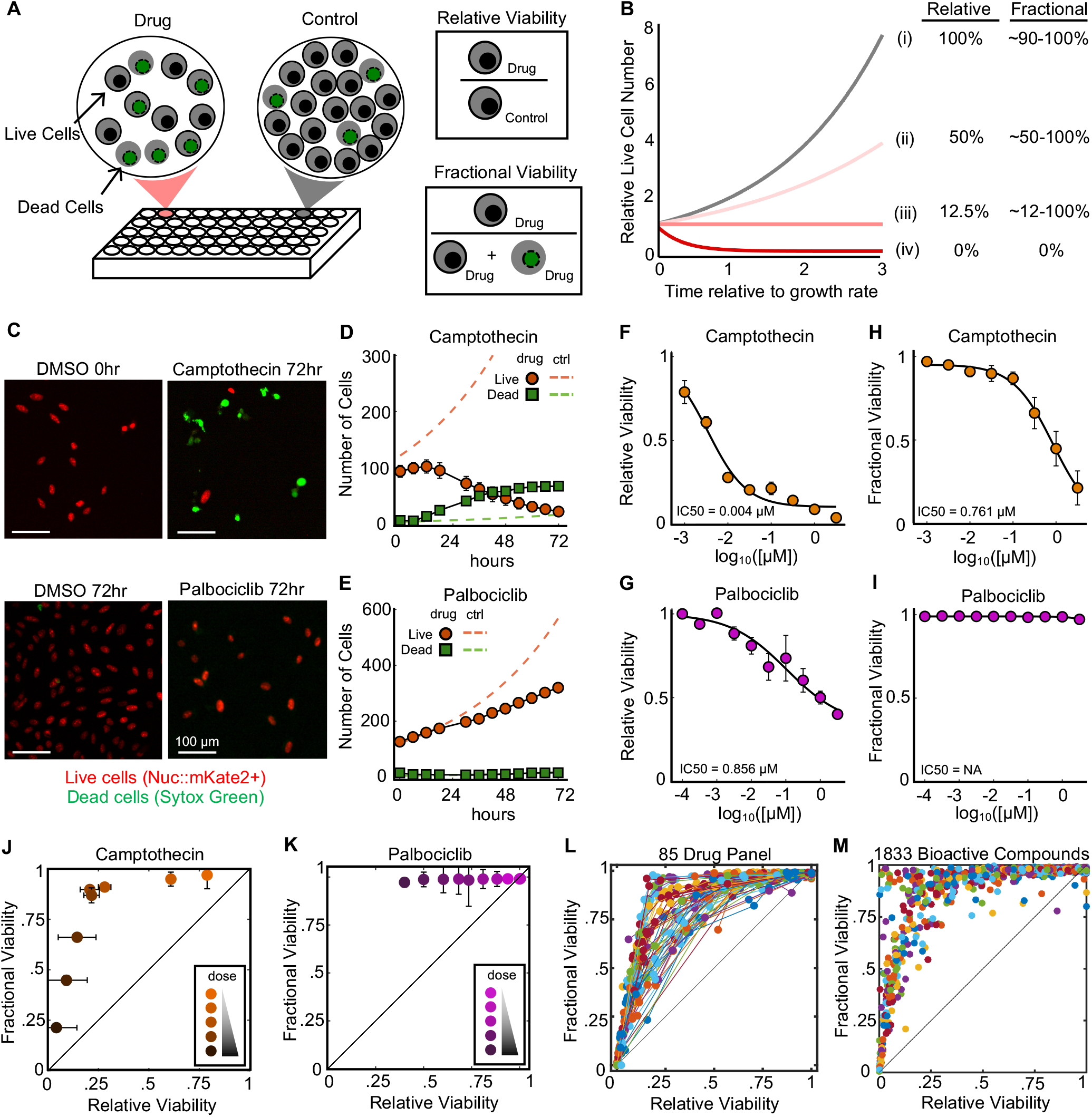
RV and FV produce largely unrelated insights into drug response. **(A)** Schematic defining common ways to quantify drug responses: Fractional Viability (FV) and Relative Viability (RV). **(B)** Simulated data of drug response over time for (i) untreated, (ii and iii) partially cytostatic/cytotoxic, and (iv) fully cytotoxic conditions. (C-I) STACK assay to measure RV and FV. U2OS-Nuc::mKate2+ cells treated with drug in the presence of SYTOX Green. **(C)** Representative images from cells treated with either DMSO, 3.16 μM Camptothecin, or 1 μM Palbociclib. **(D-E)** Quantified live and dead cell counts over time for cells as in (C). See also Supplemental Figure 1. **(F-G)** RV dose-response functions for Cam. (F) or Palbo. (G). **(H-I)** FV dose-response functions for Cam. (H) or Palbo. (I). **(J-L)** Relationship between RV and FV at all doses for Cam. (K), Palbo. (L), or 85 cell death/growth targeting drugs. See also Supplemental Figure 2. Dots for a given drug represent different doses. (M) RV vs. FV for 1833 Bioactive Compounds, each tested at 5 µM. Data in (M) are from Forcina et al.

An alternative measure of drug sensitivity does exist, in which a drug response is quantified as the fractional proportion of live and dead cells in the drug treated population (Figure 1A). This metric is variably called “lethal fraction” (or its inverse, “viable fraction”), “percent of cells”, or “percent cell death”, etc. (hereafter referred to as Fractional Viability, or FV). In contrast to RV, FV provides direct insights into the degree of cell death in a population. Additionally, Fractional Viability calculations do not require comparison between treated and untreated groups, which minimizes issues associated with plating bias, a common issue in multi-well assays (Lachmann et al., 2016). In spite of these benefits, Fractional Viability is less commonly used, because this measure generally requires extra measurements or the use of an experimental platform that provides single cell data, such as in flow cytometry based evaluation of apoptosis, or quantitative microscopy (Albeck et al., 2008; Forcina et al., 2017).

Relative and fractional measures of drug response are often used interchangeably, in spite of the fact that these are clearly different metrics (Méry et al., 2017; Riss et al., 2019). In this study, we explored the relationship between these two common measures of drug sensitivity. We find that RV and FV score unique and largely unrelated properties of a drug response. RV accurately reports the cell population size but not the degree of cell killing. Alternatively, FV exclusively reports drug-induced cell death, but does not reveal any insight into the size of the surviving population. By directly comparing relative and fractional drug responses, we find that at any given dose, most drugs induce a coincident decrease in the cell growth rate and an increase in the cell death rate. Furthermore, when evaluating across a large panel of drugs, we find a non-uniform relationship between growth inhibition and cell death, spanning the entire continuum of possible behaviors. We find that the relative proportion of drug-induced growth inhibition and cell death varies by drug, by dose, and by genotype. Furthermore, these features are not captured by traditional pharmaco-metrics such as the EC50/IC50. We develop a new quantitative analysis platform, called Drug GRADE (<u>G</u>rowth <u>R</u>ate <u>A</u>djusted <u>DE</u>ath), that captures the timing and relative magnitude of growth inhibition versus cell death. Evaluation of drug GRADE improves the ability to resolve cancer subtype-drug response relationships. Taken together, this study highlights the complex and non-uniform relationship between cell growth and cell death, and provides a new analytical framework for understanding these relationships.

## RESULTS

### Relative Viability and Fractional Viability produce largely unrelated insights about drug response

In an effort to gain deeper insights into the mechanisms of action for common anti-cancer drugs, we began by exploring the relationship between two common measures of drug response: relative viability (RV) and fractional viability (FV) (Figure 1A). A critical difference between these two measures is that RV is focused entirely on the live cell population across two conditions, whereas FV includes both live and dead cells, but only in the drug treated condition. Additionally, because RV uses an untreated control as a reference point, this measure generally cannot distinguish between responses that are due to growth inhibition versus those that are due to cell death (Hafner et al., 2016). Likewise, while decreased FV must require some degree of cell death, it is generally unclear if death occurs in a growing or growth inhibited/arrested population. Thus, while RV and FV should be correlated if not identical at extremely strong or weak response levels, the theoretical relationship between these numbers is unclear, particularly at intermediate levels of response (Figure 1B). We reasoned that exploring the relationship between RV and FV in detail might reveal hidden principles of drug sensitivity that are not captured using traditional measures. We evaluated drug responses in U2OS cells using the STACK assay, a quantitative live cell microscopy assay that measures both live and dead cells and has equal sensitivity in quantifying RV and FV (Forcina et al., 2017). We began by investigating RV and FV responses to two drugs: camptothecin, a topoisomerase I inhibitor and potent apoptotic agent, and palbociclib, CDK4/6 inhibitor that primarily induces growth arrest without inducing any cell death (Hafner et al., 2019). As expected, camptothecin induced high levels of cell death, whereas palbociclib strongly inhibited growth of the population without causing any cell death (Figure 1C-E and Supplemental Figure 1).

To characterize the relationship between RV and FV responses, we profiled each drug using an eight-point half-log dose titration. From these data, we calculated both RV and FV metrics at the assay endpoint (Figure 1F-I). A direct comparison of RV and FV for camptothecin revealed a discontinuous relationship featuring two, clearly distinct, dose-dependent behaviors (Figure 1J). In the first phase (low doses, which accounts for the majority of the RV scale), relative viability is strongly decreased in a dose dependent manner while only modestly affecting fractional viability. In the second phase (higher doses), fractional viability decreases sharply while relative viability is only modestly affected (Figure 1J). These two phases reflect a decrease in growth rate with minimal cell killing at low doses, followed by an increase in death rate, that occurs at high doses and only in growth arrested cells (Supplemental Figure 1A). Alternatively, for palbociclib which does not kill any cells, only the first of these two phases was observed (Figure 1K and Supplemental Figure 1B-C).

To determine if this bi-phasic response was a common behavior of many drugs or drug classes, we tested full dose-response profiles for a panel of 85 drugs, which target a variety of different proteins controlling cell growth and/or cell death (Supplemental Table 2). For all of these drugs, RV and FV responses were not well correlated, but the degree of correlation between these two metrics varied by drug (Figure 1L and Supplemental Figure 2). For some compounds we observed a bi-phasic dose response similar to camptothecin, characterized by two linear but discontinuous phases, with death occurring only following full growth arrest. For most drugs, however, these two phases were more mixed, and doses were found in which the RV and FV values report intermediate levels of growth inhibition and cell death. To supplement these data, we also reanalyzed a large publicly available dataset of 1833 bioactive compounds that were previously tested using the STACK assay (Forcina et al., 2017). The overall profile of responses across these diverse compounds also highlights a spectrum of behaviors, rather than exclusively bi-phasic responses. Thus, these data demonstrate that relative and fractional measures of drug response are not interchangeable, and highlight the lack of a uniform relationship between FV and RV across drugs.

**Figure 2:**
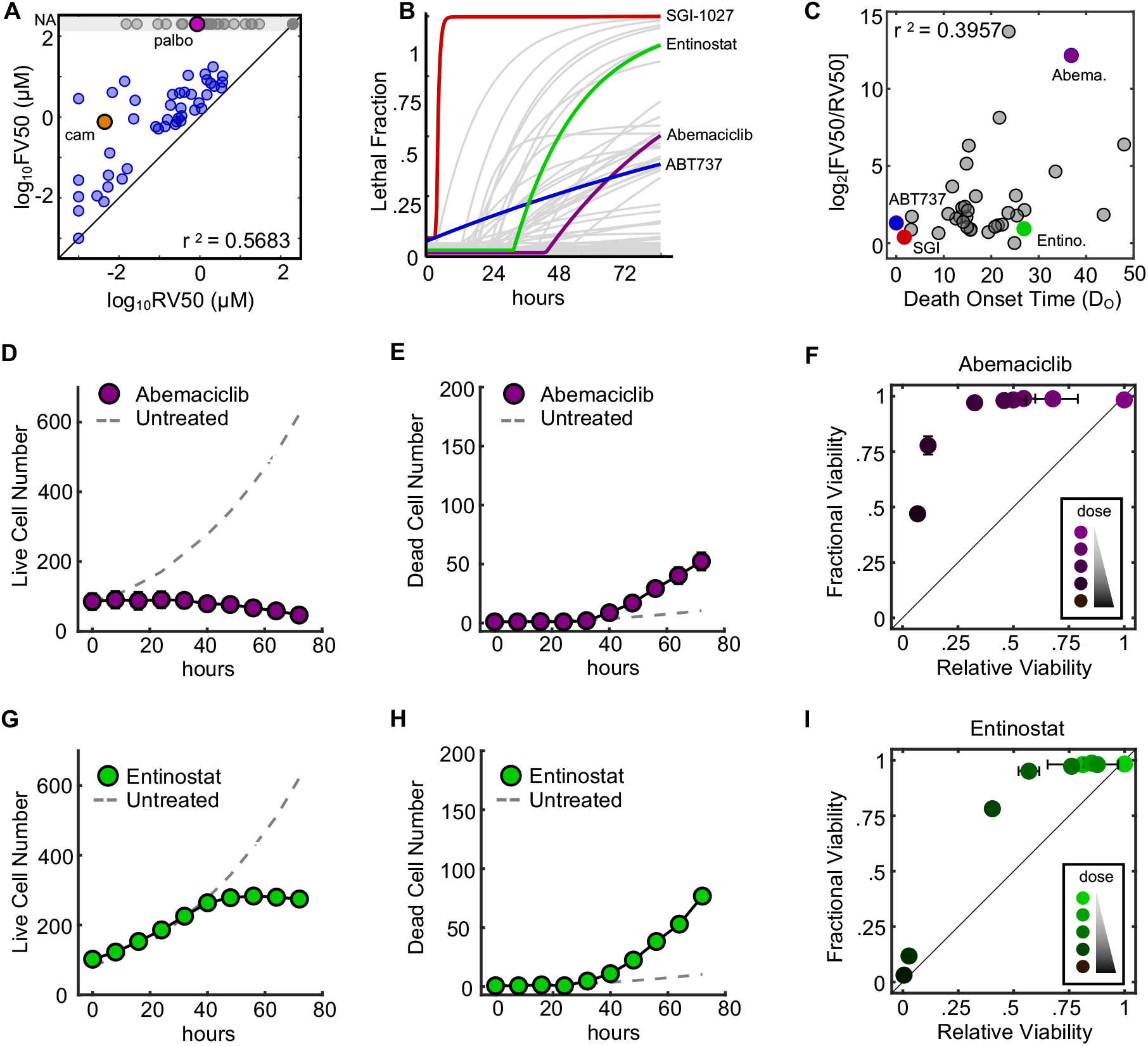
RV and FV differ due to idiosyncrasies in the strength and relative timing of drug-induced growth arrest versus cell death. **(A)** Correlation between IC50 computed using RV (RV50) or FV (FV50). Pearson correlation coefficient shown. **(B)** Death kinetics computed for 85 cell death and growth inhibiting drugs. SGI-1027 (red), Abemaciclib (purple), ABT-737 (blue), and Entinostat (green) highlighted. **(C)** Correlation between death onset time (DO) and the FV50/RV50 ratio. **(D-E)** Cell numbers over time for 10 μM Abemaciclib. (D) Live cells. (E) Dead cells. **(F)** Relationship between FV and RV for a dose range of Abemaciclib (10 μM – 0 μM) at 72hr. **(G-H)** Cell numbers over time for 3.16 μM Entinostat. (G) Live cells. (H) Dead cells. **(I)** Relationship between FV and RV for a dose range of Entinostat (31.6 μM – 0 μM) at 72hr.

### Relationships between RV and FV vary due to idiosyncrasies in the strength and relative timing of drug-induced growth inhibition versus drug-induced cell death

Overall, the IC50 doses computed using RV or FV (hereafter, RV50 and FV50, respectively) were not well correlated, often differing by several orders of magnitude (Figure 1F-J and Figure 2A). The RV50 reports the dose at which the number of live cells following drug treatment is half as large as the untreated population, whereas the FV50 reports the dose at which a population is half alive and half dead (Supplemental Figure 3). Thus, these two values should be the same only in situations where death occurs in the absence of any growth rate inhibition (i.e. death in a population of cells that are growing at the normal rate). In theory, this could be achieved several ways. For instance, drugs that induce death with very fast onset may kill cells prior to any observable changes in cell growth. Indeed, the FV50 and RV50 values were very similar for particularly fast drugs, such as SGI-1027, a DNMT1 inhibitor, and ABT-737, a BH3 mimetic (Figure 2B-C). To determine if this was a general trend, we calculated the correlation between death onset time and the FV50/RV50 ratio. We found a weak trend in which the FV50 and RV50 were more similar for drugs that had earlier onset times, but the overall correlation was modest, suggesting that death onset time alone was not a particularly good predictor of the FV/RV relationship (r^2^ = 0.3957, Figure 2C).

**Figure 3:**
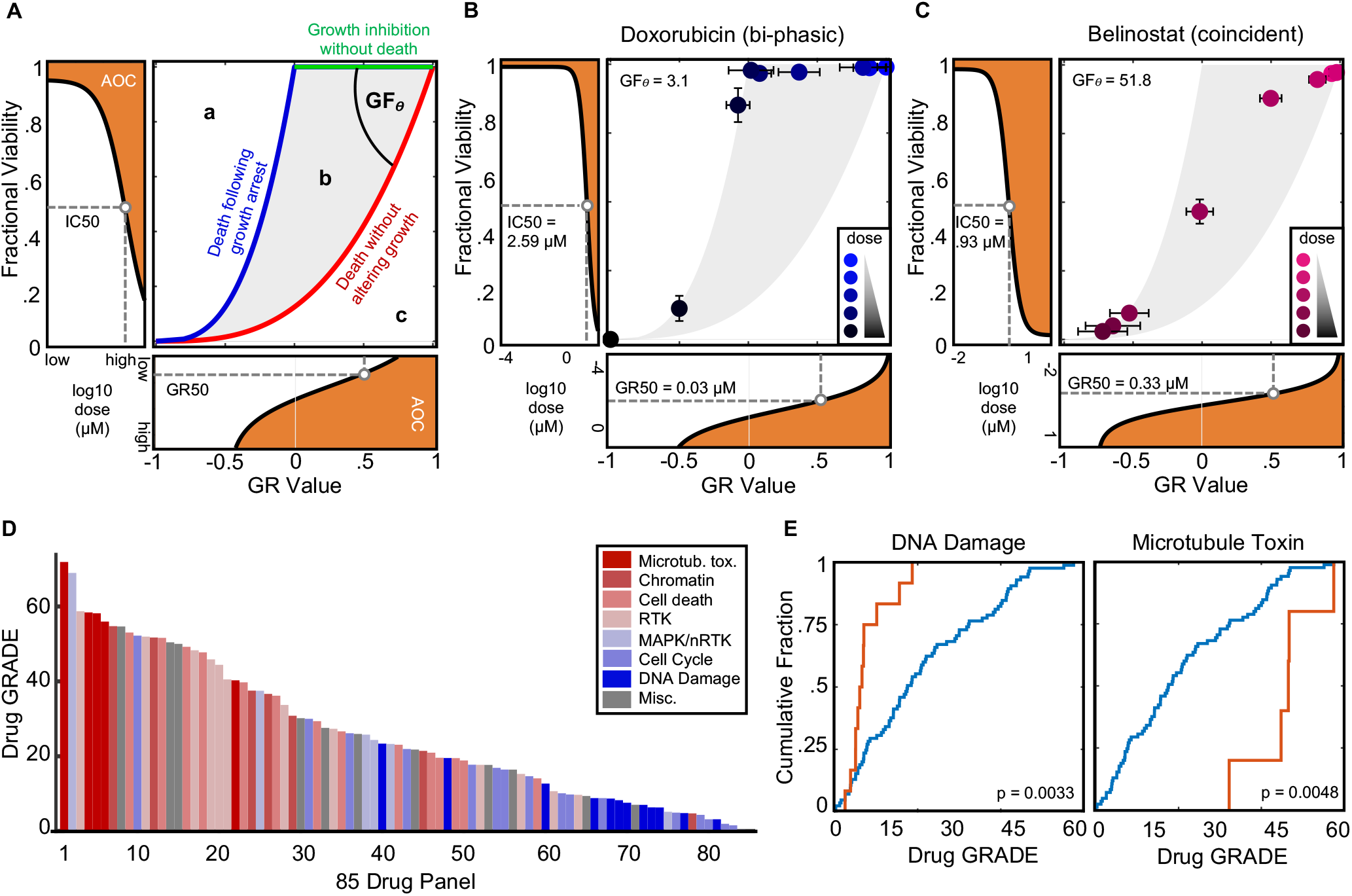
Integrative analysis of relative and fractional drug responses reveals a continuum of distinct relationships between drug-induced growth arrest and cell death. **(A)** Schematic of the GR/FV plot with reference limits shown for growth inhibition without death (green), death only following growth arrest (blue), and death without altering growth rate (red). Shaded region “b” represents intermediate states in which a drug induces some growth inhibition and some death. GR*θ* scores the relationship between FV and GR measures and reports the contribution of cell death to the observed population reduction. See also Supplemental Figure 5. **(B)** GR/FV plot for an example bi-phasic drug, doxorubicin. **(C)** GR/FV plot for Belinostat, a drug that induces coincident growth rate inhibition with cell death. **(D)** Waterfall plot of GF*θ* scores for 85 drugs tested. See also Supplemental Figure 7. **(E)** Cumulative distribution functions of GF*θ* scores for all 85 drugs (blue) or drugs in the listed class (orange). P values calculated using the a two-tailed KS test.

In theory, other mechanisms exist, in addition to death onset time, that likely account for variations between FV and RV metrics. For instance, regardless of death onset time, FV and RV values would differ if a drug induces potent growth inhibition at low, non-killing doses, as we observed for drugs that induce bi-phasic responses, such as camptothecin (Figure 1J). Likewise, even for drugs with a very late death onset time, FV and RV values should be similar if the onset time of growth inhibition is also equally late. To identify such scenarios, we focused on drugs for which the death onset time was a poor predictor of the relationship between FV and RV, such as Abemaciclib and Entintostat.

FV50 and RV50 values for the CDK4/6 inhibitor Abemaciclib were unusually varied, even for a drug with slow death onset time. Consistent with our expectations, Abemaciclib produced a distinctly bi-phasic dose response, characterized by growth inhibition at low doses, and death only at high doses. (Figure 2D-F). Furthermore, our comparisons of RV and FV values over time, rather than across doses, revealed that Abemaciclib induces death only following a prolonged period of growth arrest (Supplemental Figure 4). A discontinuous bi-phasic responses for Abemaciclib was also recently reported to occur due to its off-target activity against CDK2 (Hafner et al., 2019).

**Figure 4:**
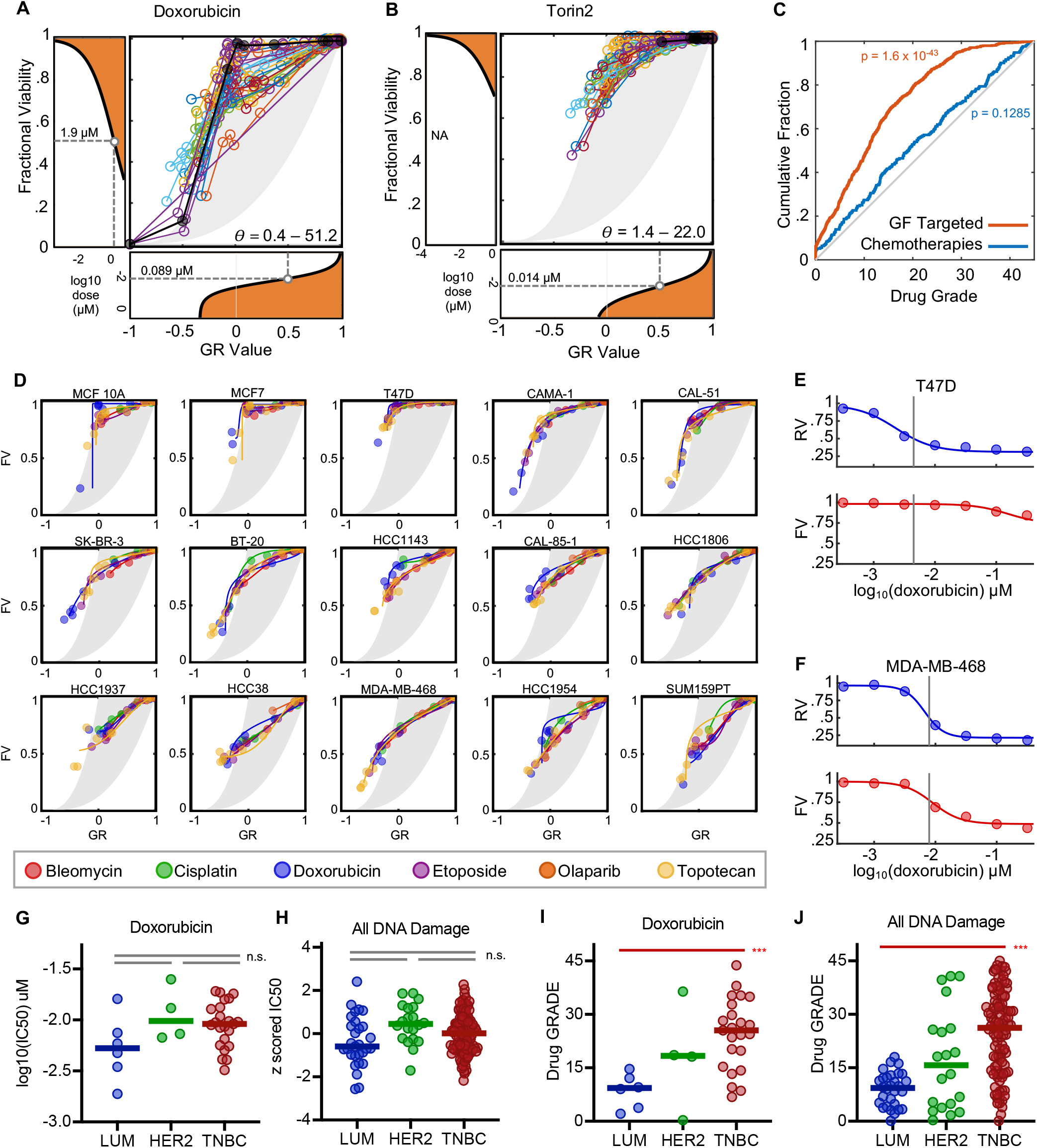
Drug GRADE captures subtype dependent differences in drug sensitivity that are not captured using traditional pharmaco-metrics. **(A-B)** GR/FV plots for Doxorubicin (A) or Torin 2 (B) for U2OS (black) and 35 different cell lines from the LINCS dataset. Range of GF*θ* scores across all cell lines shown. GR and FV Dose curves are for the mean responses across all cell lines. **(C)** CDF plot of GRq scores for cytotoxic chemotherapies or growth factor targeted therapies. P-value from KS test shown for deviation from random scores. **(D)** GR/FV plots for 6 DNA damaging drugs across 15 breast cancer cell lines from LINCS. See also Supplemental Figure 8. **(E-F)** RV and FV dose responses shown for doxorubicin in T47D (E) or MDA-MB-468 (F). Traditional IC50 (i.e. RV50) highlighted with grey bar. **(G-H)** Traditional IC50s for Doxorubicin (G) or all DNA damaging drugs (H) across 36 LINCS cells. Data are separated by sublass: luminal (LUM), HER2 over-expressing (HER2), or triple-negative (TNBC). (I-J) Drug GRADE for Doxorubicin (G) or all DNA damaging drugs (H) across 36 LINCS cells. Data are separated as in (G) and (H). For panels G-J, t-test p-values are shown for comparison of TNBC to LUM. All other comparisons are not significant.

Alternatively, the HDAC inhibitor Entinostat induced death with a delayed onset time of approximately 30 hours after drug exposure, but nonetheless FV and RV values were well correlated (Figure 2B). For this drug, kinetic analysis revealed that Entinostat treated cells proliferate at precisely the untreated growth rate for approximately 30 hours, such that the onset time of growth inhibition is equally delayed and similar to the onset time of cell death (2G-I). Thus, even for a relatively small set of drugs, these data highlight the lack of a singular “rule” describing the relationship between FV and RV values. The relationship between FV and RV depends on a combination of features, including the death onset time and whether cell death is occurring in a proliferating or growth arrested population. Taken together, these data underscore the fact that common pharmaco-metrics derived from FV or RV fail to capture the relationship between drug-induced changes in growth versus cell death.

### Integrative analysis of relative and fraction drug responses captures distinct drug class specific relationships between growth inhibition and cell death

Relative viability measures different aspects of a drug response than Fractional Viability. Because a simple rule could not be identified for predicting one from the other, we next asked what might be learned by quantitatively exploring the relationship between these metrics. We began by simulating RV and FV values for theoretical drug responses, using all possible combinations of fractional growth inhibition and fractional cell death in different proportions (Supplemental Figure 5A-B). These simulations revealed an area of possible responses with boundaries representing three distinct response scenarios: growth arrest without any cell death (green line, top, Supplemental Figure 5C), cell death within a population of normally growing cells (red line, right, Supplemental Figure 5C), and a discontinuous bi-phasic response characterized by growth arrest at low doses, followed by cell death only in growth arrested cells (blue, top and left, Supplemental Figure 5C).

The size and shape of this region varies dramatically depending on the length of the assay and the growth rate assumed in the simulation. Thus, to stabilize these relationships, we also simulated drug responses using the normalized Growth Rate Inhibition (GR) value. GR values are similar to RV in that both are derived from measurements of live cells in drug treated and untreated conditions. A critical difference, however, is that the GR value scores a drug response based on a comparison of population growth rates in the presence and absence of drug, rather than scoring changes in population size as in RV (Hafner et al., 2016). Thus, GR corrects for artifactual differences in drug sensitivity that might be caused by differences in assay length between experiments or differences in proliferation rate between cell types. Importantly, a comparison of simulated FV and GR values revealed a region of possible relationships defined by the same boundaries seen for FV vs. RV comparisons (Figure 3A and Supplemental Figure 5D).

For both FV vs. GR and FV vs. RV comparisons, the area between the observed limits represents drug responses that feature both some growth inhibition and some cell death at varied proportions. Importantly, from the simulated data, any point within this bounded space can be attributed to a specific proportion of fractional growth inhibition and cell death (Supplemental Figure 5). The regions outside of the bounded area represent responses that, while conceptually possible, are not observed in our simulated responses. Region “a” to the left of the bounded area would include drug responses in which the population size is decreased in excess of the measured number of dead cells (Supplemental Figure 5C). This may be observed for some types of cell death, such as entosis (Overholtzer et al., 2007), and for technical reasons related to assay precision and/or the relative sensitivity of live cell and dead cell measurements. Region “c”, to the right of the bounded area, includes responses in which the degree of cell death is compensated for by a drug-induced increase in the growth rate.

Importantly, although responses in region “a” and “c” are possible in theory, these are never observed in our experimental data, which for all 85 drugs profiled fell entirely within the bounds represented by region “b” (Supplementary Figure 6). Interestingly, several drug’s responses fell precisely at the left most boundary, represented by bi-phasic dose response profiles, including Abemaciclib and most DNA damaging chemotherapeutics (Figure 3B and Supplemental Figure 6). Most drug responses, however, were characterized by GR and FV values that reveal partial growth suppression that occurs coincidentally with partial cell death at different proportions for each drug (Figure 3C).

To characterize these drug-specific differences in the GR and FV values, we calculated the <u>G</u>rowth <u>R</u>ate <u>A</u>djusted <u>De</u>ath percentage (GRADE). The drug GRADE reports the proportion of an observed reduction in population size that is due to cell death. We calculated the drug GRADE using the angle formed between the observed FV/GR data and a perfectly bi-phasic drug response (Figure 3A, GF_θ_). This angle is further rescaled relative the minimum and maximum angles possible within our simulated data, such that drug GRADEs scale from 0 – 100, with 100 reporting that the observed response was entirely due to cell death and a GRADE of 0 reporting that the observed response was entirely due to growth inhibition. Inspecting drug GRADE for the 85 drugs that we profiled revealed a continuous distribution of values, further demonstrating the unique drug-specific relationship between growth inhibition and cell death (Figure 3D). Nonetheless, similarities were also observed between drugs of a given class. For instance, DNA damaging chemotherapeutics were enriched for very small drug GRADEs, indicating that for these drugs, the population reduction at IC50 doses is generally due to growth inhibition, rather than cell death. Alternatively, microtubule toxins tended to have large drug GRADEs indicating potent killing at IC50 doses (Figure 3E and Supplemental Figure 7). Importantly, drug GRADE was not correlated with traditional pharmaco-metrics, such as the IC50, EC50, or Emax (Supplemental Figure 8). Thus, while traditional pharmaco-metrics report insights into drug affinity, potency, or efficacy, drug GRADE provides a unique insight into the mechanism of population reduction.

### Drug GRADE captures subtype dependent differences in drug sensitivity that are not captured using traditional pharmaco-metrics

Drug potency and drug efficacy are known to vary in a genotype and cancer subtype dependent manner. It was unclear if drug GRADEs are stable features of a given drug, or if these would also vary for a given drug across genotypes or cancer subtypes. To explore this question, we analyzed a publicly available dataset collected by the LINCS consortium, which contained 34 drugs tested across 35 breast cancer cell lines, with the data collected in a manner that would allow both GR and FV calculations (Hafner et al., 2019). For essentially all drugs, we found striking differences in drug GRADE across the cell lines (Supplemental Figure 9). For instance, doxorubicin, a topoisomerase II inhibitor that is commonly used in the treatment of breast cancer, produced a bi-phasic dose response in U2OS cells, characterized by killing only at high doses, and only following full growth arrest (GF_θ_ = 3.1; Figure 4A). In the LINCS breast cancer cell lines, however, the GF_θ_ values ranged from 0 – 51, revealing substantial variation in the degree of cell killing at IC50 doses (Figure 4A). Variation in drug GRADE was observed for all drugs, including targeted agents like Torin2 (Figure 4B). Cell cycle and growth factor targeted therapies were skewed towards smaller GF_θ_ values, consistent with the notion that these drugs primarily induce growth inhibition, rather than cell death. Interestingly, cytotoxic chemotherapies, which can induce both growth inhibition and cell death, had a nearly random distribution of drug GRADEs across the genotypes studied (Figure 4C).

For cytotoxic chemotherapies, the observed variance in drug GRADE across cell lines might suggest that drug GRADE can capture genotype specific differences in drug response. An alternative explanation could be that the relationship between drug-induced growth arrest and drug-induced cell death is not determined by the drug, but instead is either stochastic or subject to strong environmental/context dependent regulation. To distinguish between these possibilities, we investigated variation within GF_θ_ values for drugs that share a common mechanism within each individual cell line. The LINCS dataset includes 6 different drugs that act by causing DNA damage, and 10 drugs annotated as PI3K/mTOR inhibitors (Supplemental Table 2). For any one of these drugs, significant variation was observed in drug GRADE across the LINCS cell lines (Figure 4A-B, and Supplemental Figure 9). In contrast, within any given cell line, drugs of a shared class produced strikingly similar drug GRADEs (Figure 4D and Supplemental Figure 10). Similar drug GRADEs were observed even for the DNA damage drug class, which included drugs that induce DNA damage using a variety of different molecular mechanisms and through unrelated drug binding targets. These data suggest that the variation observed for drug GRADE is related to the specific ways in which a given cell/cell type responds to a class of drugs.

The variations that are uncovered by drug GRADE reveal important differences in the underlying drug response. For instance, DNA damaging agents resulted in bi-phasic dose responses in T47D, a luminal ER+ breast cancer cell line (mean GF_θ_ = 4.1; Figure 4D). In contrast, these drugs consistently resulted in coincident growth inhibition and cell death in MDA-MB-468, a basal triple-negative breast cancer (TNBC) cell line (mean GF_θ_ = 31.9; Figure 4D). This distinction reveals that the traditional IC50 (IC50 calculated from RV, RV50) captures a partially growth suppressing dose in T47D, but the same pharmacological value captures a potent killing dose in MDA-MB-468 (Figure 4E-F). Furthermore, while the IC50 values are similar and not statistically distinguishable for most DNA damaging drugs in these two cell lines, they are generally lower in T47D when compared to MDA-MB-468, and generally lower in luminal cells when compared to TNBCs (Figure 4G-H). Thus, from the IC50 data alone, one might predict either equal chemosensitivity among breast cancer subclasses or that luminal breast cancer cells are more chemosensitivity than TNBC. These conclusions would be inconsistent with established clinical data, as TNBCs are well-validated to be more chemosensitive than other breast cancer subtypes (Carey et al., 2007). Importantly, while the IC50 fails to capture subtype-specific differences in chemosensitivity, drug GRADE identifies significant differences between breast cancer subtypes. DNA damaging drugs in TNBC have significantly higher drug GRADEs than in other breast cancer subtypes, revealing that DNA damaging chemotherapies induce greater levels of cell death in TNBC than in other breast cancer subtypes (Figure 4I-J). Taken together, these data highlight that drug GRADE captures critical differences in drug response that are not captured by traditional pharmaco-metrics.

## DISCUSSION

Recent studies that have revealed that differences in the cell growth rate are a confounding factor in the measurement of the effect of anti-cancer therapies (Hafner et al., 2016; Harris et al., 2016). These studies were a major step forward in analysis methods and have provided much needed clarity into drug-induced changes in population size. The strategy we use here builds upon these prior works, and in fact, uses the GR metric as one of the two key analysis features. A clear distinction, however, is that our approach integrates an independent measurement of dead cells and drug-induced Fractional Viability. We find that the integrated analysis of population growth (through GR) and fractional killing (through FV) reveals drug- and cancer subtype-specific features of a drug response that are not captured using either of these values alone, or when using any traditional pharmaco-metrics.

The most common measures of drug response are derived exclusively from measurements of live cells. Using these measurements to infer of the degree of death requires some assumption to be made about the relationship between drug-induced growth inhibition and cell death. For instance, a common assumption is that cell death occurs only in growth arrested cells. A central finding from our study is that the relationship between drug-induced growth inhibition and cell death varies substantially across drugs in a continuous manner. Also, for a given drug or drug class, drug GRADE varied substantially across cancer subtypes. Thus, in the absence of direct measurement of FV and RV-type responses, any assumption made regarding the relationship between growth inhibition and cell death is certain to be wrong in most situations.

Of note, the sign of the GR scale is generally interpreted as revealing the response phenotype, with positive GR values interpreted as partial growth inhibition whereas negative values are interpreted as cell death (more formally interpreted as a negative growth rate, or death rate). Although it must be true that negative GR values report drug-induced cell death, notably, positive GR values do not necessarily report the lack of cell death. This was clearly demonstrated in theory in the original description of the GR value (Hafner et al., 2016), and our analysis reveals that for most drugs, significant levels of death are observed in the positive portion of the GR scale. These phenotypes generally resulted from intermediate levels of cell death occurring in a population of cells that continue to proliferate. Thus, while the GR value unambiguously reports the net population growth rate in a manner that distinguishes between an increasing and a decreasing population size, whether or not a drug induces significant killing requires additional measurements. The strategy we describe in this study clarifies this issue and our data show that GR and FV values provide complementary insights into the nature of a drug response.

One limitation of the analysis method we propose is that it cannot be used in conjunction with many common drug-response assays that exclusively measure live cells (CellTiter-Glo, MTT, alomar blue, colony formation, etc.). Our approach should be amenable to any assay that develops single cell data for live and dead cells, such as flow cytometry, histology, or the microscopy-based STACK analysis used in this study. Additionally, we recently developed a high-throughput fluorescent plate reader-based strategy for inferring live cell counts using only a direct measurement of dead cells (Richards et al., 2019). Thus, if only live or dead cells can be counted, our data suggest that measurement of dead cells would be preferable, as live cells can be accurately inferred using modest experimental and computational adjustments (Richards et al., 2019).

Drug response assays are common to many sectors of biomedical research, and a common practice is to summarize drug responses using measures such as the IC50, EC50, or Emax. These metrics are used to compare across drugs or to compare drug responses across biological scenarios. In many situations, such as oncology, a critical question generally remains unanswered by these metrics: does the drug actively kill cells or just result in growth inhibition? This is an important distinction, as growth inhibition is not likely to be a durable response, particularly when considering the rapid clearance of most chemotherapeutics due to drug metabolism and excretion. In current approaches, a common strategy to determine if an observed response is due to cell death or growth inhibition is to use relative viability to characterize drug potency/efficacy. These measures are then complemented with a more specific measure of cell death, in order to determine if the observed response was caused by growth arrest or cell death. Our study reveals a flaw in this line of thinking, that the response was necessarily “either/or” and not “both”. We find that most drugs achieve their effects through some combination of growth inhibition and cell death, but the relative proportions of these effects vary by drug and by across different cancer subtypes. Clarifying these relationships should improve our ability to accurately evaluate drug responses and how these responses vary across drugs or across biological contexts.

## METHODS

### Cell lines and reagents

mKate2 expressing U2OS cells were generated as previously described (Richards et al., 2019). Cells were grown in Dulbecco’s modified eagles medium (DMEM) supplemented with 10% FBS, 2 mM glutamine, and penicillin/streptomycin. Chemicals and drugs were purchased from either Apex Biologics/ApexBio, Selleck Chemicals, or Sigma. For detailed information see Supplemental Table 1.

### Quantitative live cell imaging and image analysis

U2OS::mkate2+ cells were grown in 10cm dishes (Fisher Scientific, Cat #: FB012924). Prior to drug treatment (“ Day −1”), cells were trypsinized, counted using a hemacytometer, and plated at a concentration of 2500 cells per 90 µL of media. Experiments were performed in 96-well black-sided optical bottom plates (Corning, Cat #: 3904). Following overnight incubation, drugs were added in growth media containing 500nM SYTOX Green (10 µL volume; final concentration of 50 nM SYTOX in the well). Images was collected using the STACK assay (Forcina et al., 2017). Briefly, images were acquired using the IncuCyte S3 with settings for the green channel: ex: 460 ± 20; em: 524 ± 20; acquisition time: 300ms; and red channel: ex:585 ± 20; em: 635 ± 70; acquisition time: 400ms. Data were acquired either every 6-8 hours for 72 hours, or only at 72 hours when kinetic analysis was not needed.

Live and dead cells were counted using the IncuCyte built-in software. Cell counting parameters were empirically determined using untreated cells and a subset of cytotoxic compounds. Analysis settings for SYTOX Green+ objects were: Top-Hat segmentation; Radius(μm) between 50 and 100; Threshold(GCU) between 5 and 10; Edge split on; Edge sensitivity between −25 and −45; Filter area min between 20 and 55; Filter area max between 2600 and 3000; Max eccentricity between 0.90 and 0.95. Analysis settings for mkate2+ objects were: Top-hat segmentation, Radius(μm) between 100 and 110; Threshold(GCU) between 0.8 and 1; Edge split on; Edge sensitivity between −45 and −35; Filter area(μm^2^) max between 100 and 110; Filter area(μm^2^) max between 2600 and 3000. The counts per well for the Sytox+ and mkate2+ objects were exported to excel and loaded into MATLAB for further analysis. For some experiments that did not require kinetic analysis, images were acquired using a EVOS FL Auto automated microscope. For images obtained using the EVOS microscope, the images were analyzed using custom MATLAB scripts, available upon request.

### Calculation of Drug GRADE

Live cell and dead cell data generated from microscopy were used to calculate “fractional viability” (live cells divided by total cells; FV). Growth rate inhibition metrics (GR) were calculated as described (Hafner et al., 2016). See also Supplemental Figure 5. To calculate drug GRADE, we focused on all doses of a given drug that are less than or equal to the GR50 dose. Our experimental and simulated data show that the relationship between FV and GR is roughly linear for GR values between 0.5 – 1. Thus, for these doses the relationship between GR and FV were fit to a linear function. Drug GRADE was determined using the following equation:

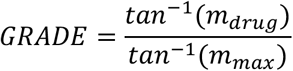

where tan^−1^ is the inverse tangent (‘atan’ function in MATLAB), *m*_*drug*_ is the slope of the linear fit relationship between FV and GR for doses of GR less than or equal to the GR50 dose, and *m*_*max*_ is the maximum slope observed over the same range of GR values, given the assumption that the observed response was entirely due to cell death, without any drug-induced growth slowing. The maximum possible slope was determined from simulated experiments as described in Supplemental Figure 5. Thus, drug GRADE reports as a percentage the contribution of cell death to the observed response at IC50 dose.

### Data analysis and statistics

Data from IncuCyte imaging was analyzed using the IncuCyte built-in software, as described above. All other data were analyzed in MATLAB. GR transformations of RV data were performed as described previously (Hafner et al., 2016). Dose-response functions were generated using a four-parameter logistic regression as described previously [ref]. Theoretical and experimental growth curves and associated growth rates were calculated in MATLAB using the built-in function ‘fit’ and the growth model ‘a*2^bx^’. Death kinetic rates (DO and DR) were determined using MATLAB, as described previously (Richards et al., 2019). Statistical enrichments were determined in MATLAB using built-in functions ‘kstest2’ or ‘fishertest’ as indicated in the figure legends.

### Data and code availability

Source data collected for a panel of 85 drugs at varied doses in U2OS cells are included in Supplemental Table 2. Images and raw cell counts from images will be made available upon request. Custom MATLAB code for computing drug GRADE and generating FV/GR plots are included in Supplemental Code 1. Custom MATLAB scripts for image analysis or curve fitting will be made available upon request.

## ACKNOWLEDGEMENTS

We thank current and past members of the UMassMed PSB community for their helpful comments and critiques during the design and execution of this study. Additionally, we thank M. Hafner, S. Peyton, J. Pritchard, and M. Walhout for their helpful comments during the preparation of this manuscript. This work was supported by National Institute of General Medical Sciences of the National Institutes of Health (R01GM127559 to MJL); the American Cancer Society (RSG-17-011-01 to MJL); and a NIH/NCI training grant (Translational Cancer Biology Training Grant, T32-CA130807 to RR).

## AUTHOR CONTRIBUTIONS

This project was conceived by HRS, RR, and MJL. Drug response data were collected by HRS, RR, REF, and AJJ. Data analysis was performed by HRS, RR, MEJ, and MJL. The manuscript was written and edited by HRS, RR, and MJL.

## CONFLICTS OF INTEREST

The authors declare no conflicts of interest

## SUPPLEMENTAL FIGURE LEGENDS

**Supplemental Figure S1:**
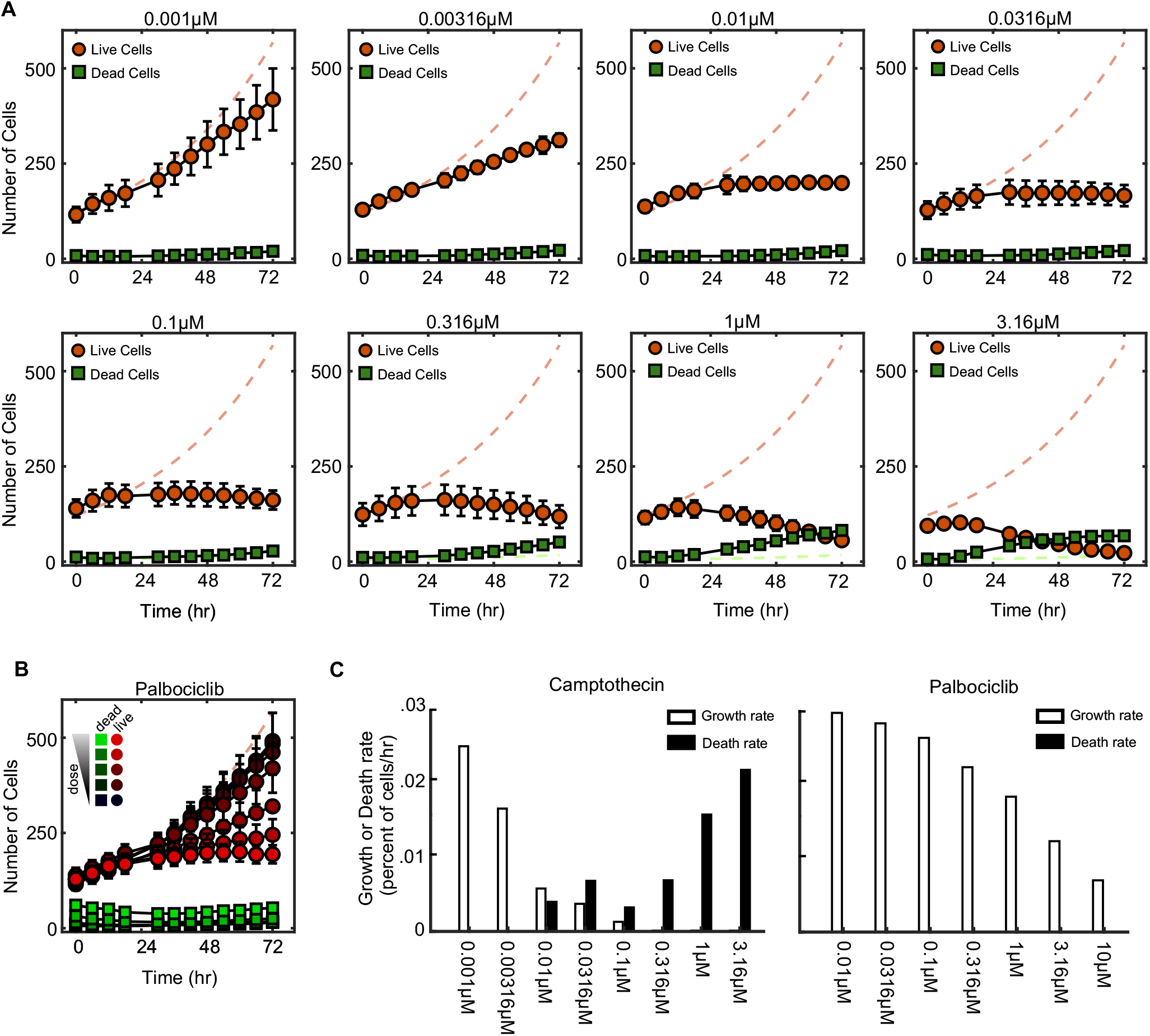
Camptothecin and palbociclib kinetic responses reveal bi-phasic dose response behavior. **A)** the number of live and dead cells present over a 72-hour time course at different concentrations of Camptothecin. Red dashed line shows the number of live cells present in the untreated condition. Green dashed line shows the number of dead cells present in the untreated condition. Dashed red and green are live and dead cell numbers for control untreated cells, respectively. **B)** The number of live and dead cells present over a 72-hour time course at different concentrations of Palbociclib (10, 3.16, 1, 0.316, 0.1, 0.0316, or 0.01 uM). Data for **A** and **B** was collected using the STACK assay. Images were acquired every 6 hours. Error bars indicated the standard deviation across four replicates. C) The growth rate and death rate was calculated using an exponential growth model or lag-exponential death model, respectively, for each dose of camptothecin and palbociclib shown in panels **A** and **B**.

**Supplemental Figure S2:**
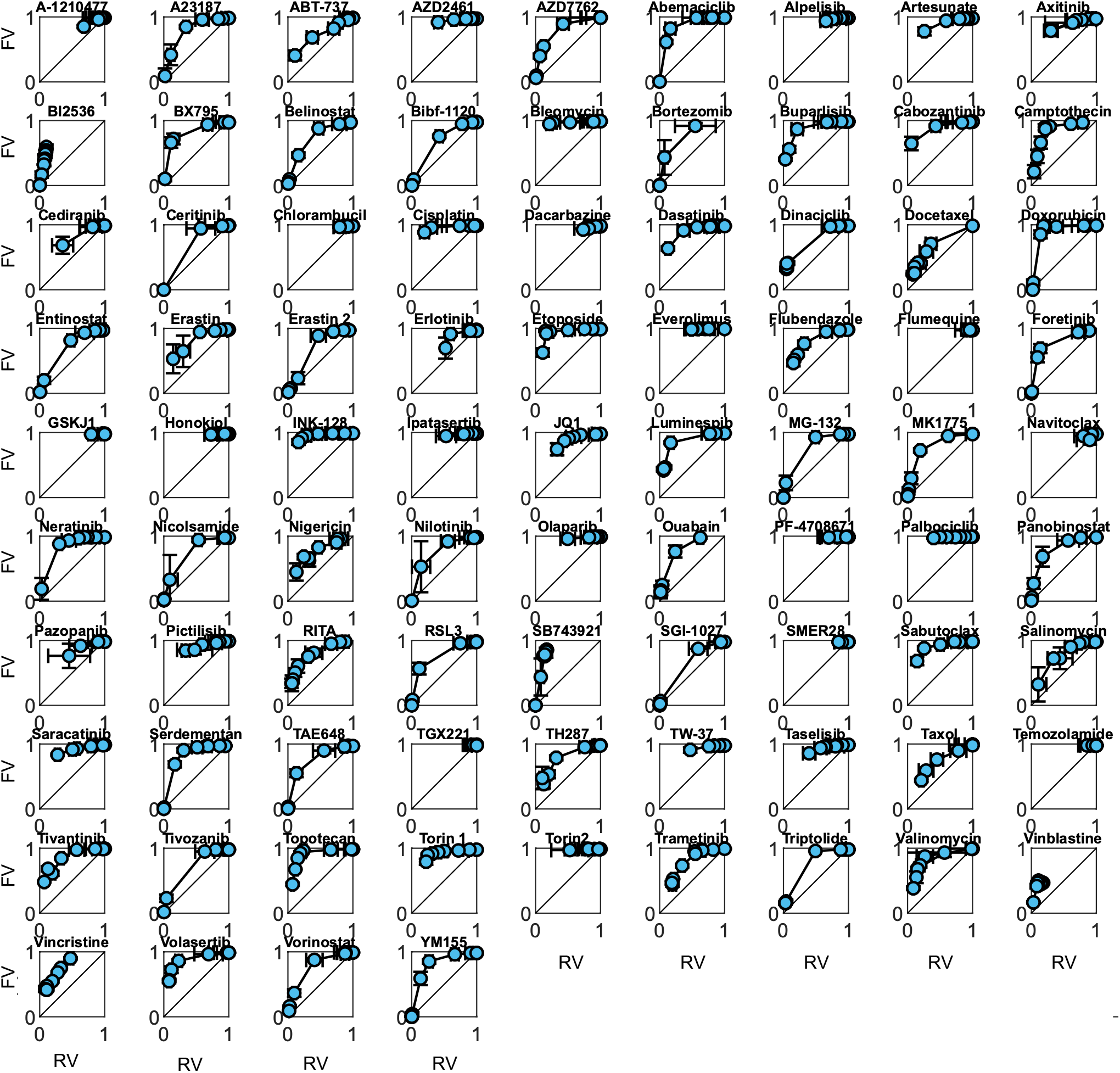
RV versus FV plots for all 85 drugs tested in U2OS. RV versus FV plots for 85 chemotherapies tested in U2OS. Error bars represent the standard deviation across 4 biological replicates.

**Figure S3:**
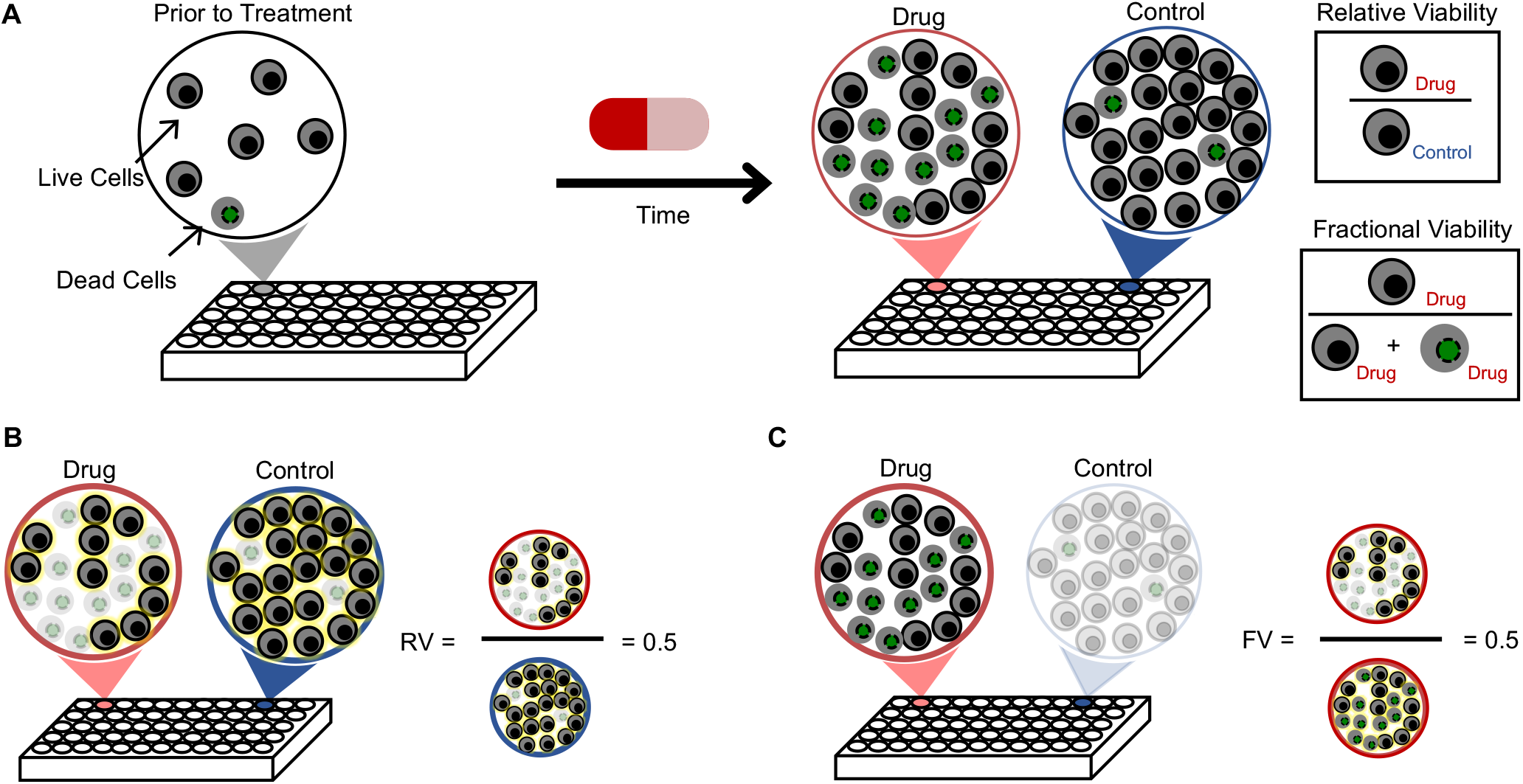
Schematic of the relationship between RV50, FV50, and GR metrics. **(A)** Schematic of a typical drug response assay in multi-well plates. Relative Viability (RV) and Fractional Viability (FV) are two common measures of response. **(B)** RV calculated for the example shown in (A). The example shown represents the IC50 of the RV measure (i.e. “RV50”), defined as the dose at which the observed number of live cells after drug exposure is half the size of the untreated population. **(C)** FV calculated for the example shown in (A). The example shown represents the IC50 of the FV measure (i.e. “FV50”), defined as the dose at which the population is half alive and half dead. Note, although both values in (B) and (C) are 0.5, these are computed from different comparisons and are defined differently.

**Figure S4:**
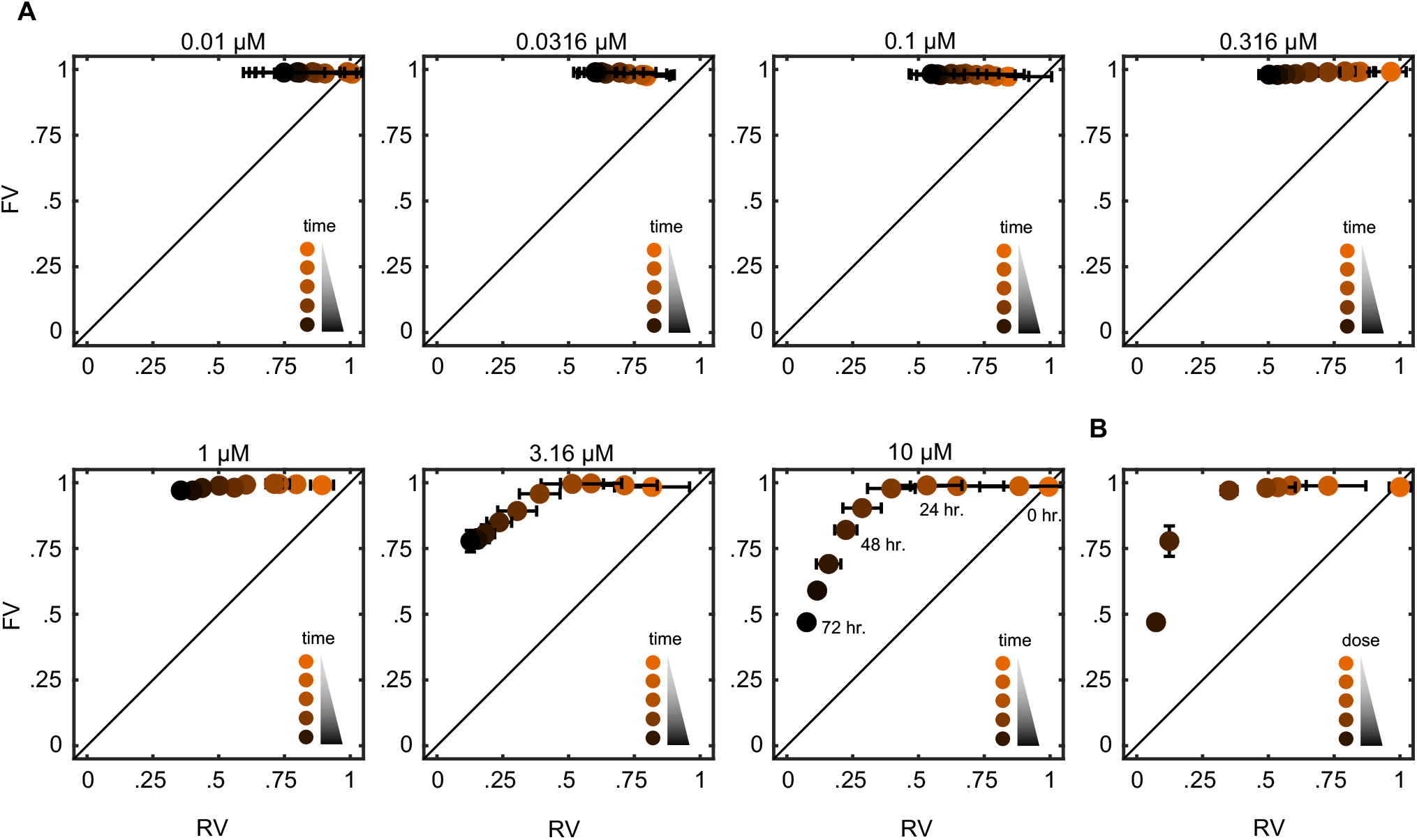
Perfectly bi-phasic transition through growth arrest before causing cell death. **A)** RV/FV plots for each dose of Abemaciclib tested over a 72-hour time course using the STACK assay. Images were acquired every 8 hours. **B)** RV versus FV plot for varied doses 72-hours after drug exposure. For the data shown in panel **A**. Error bars indicate standard deviation across two replicates.

**Figure S5:**
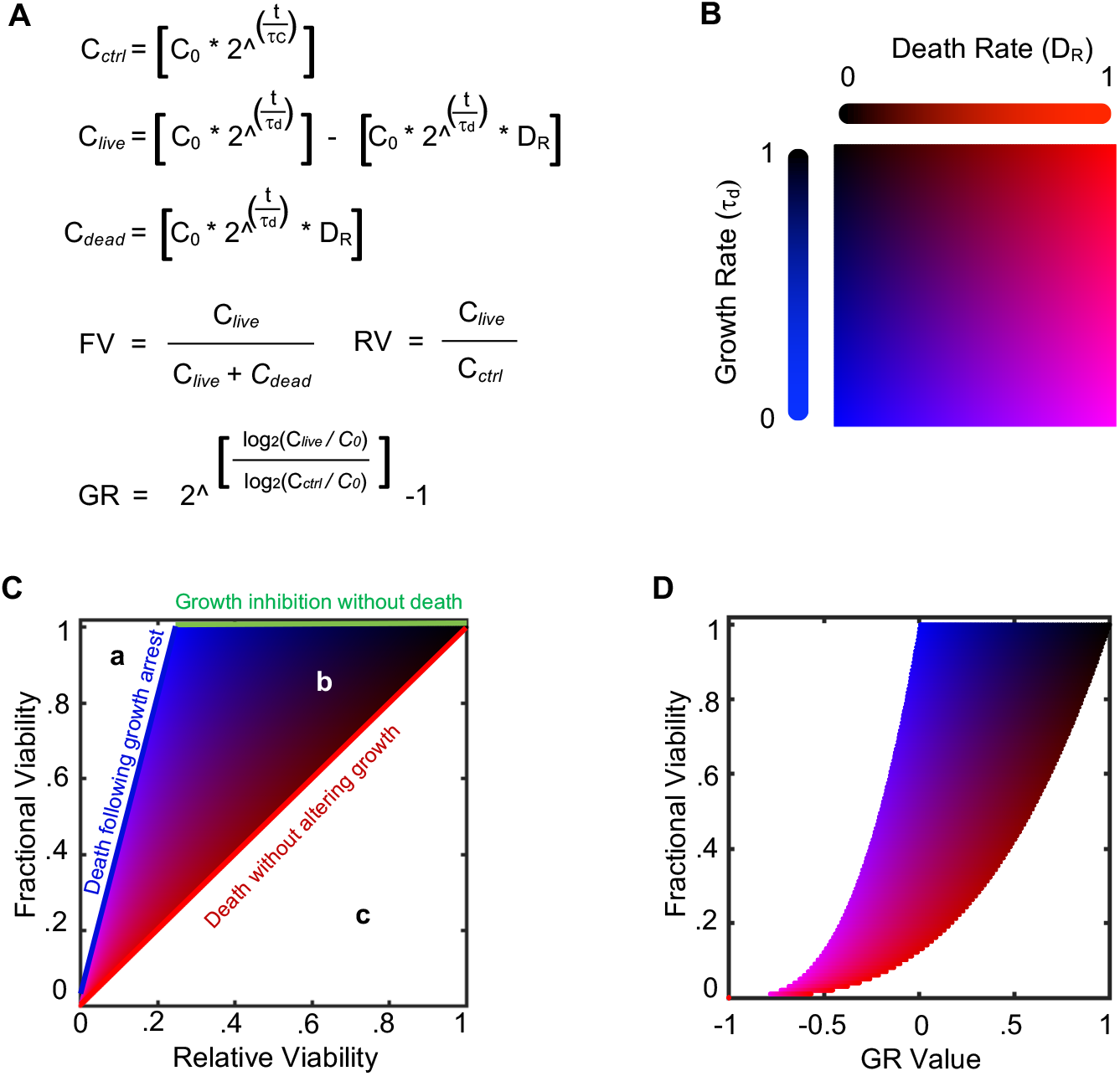
Simulations of drug-induced changes in growth rate and death rate. **(A)** Equations for live cells in control untreated condition (C_*ctrl*_), live cells in drug treated condition (C_*live*_), dead cells in drug treated condition (C_*dead*_), Fractional Viability (FV), Relative Viability (RV), and growth rate inhibition metrics (GR). C0 = initial cell number; t = assay duration; tc = growth rate of control (untreated cells); td = growth rate of drug treated cells; DR = death rate of drug treated cells. For this simulation, the death rate of control cells is presumed to be zero. **(B-D)** Simulated drug responses using all possible variations of drug-induced growth rate and death rates. **(B)** Color map of parameter values. Red increases as death rate increases. Blue increases as growth rate decreases. The scale for D_R_ is in fraction of cells per hour. The scale for Growth Rate is fraction of control untreated growth rate. **(C)** FV and RV calculated for full parameter space in (B). **(D)** FV and GR calculated for full parameter space in (B).

**Figure S6:**
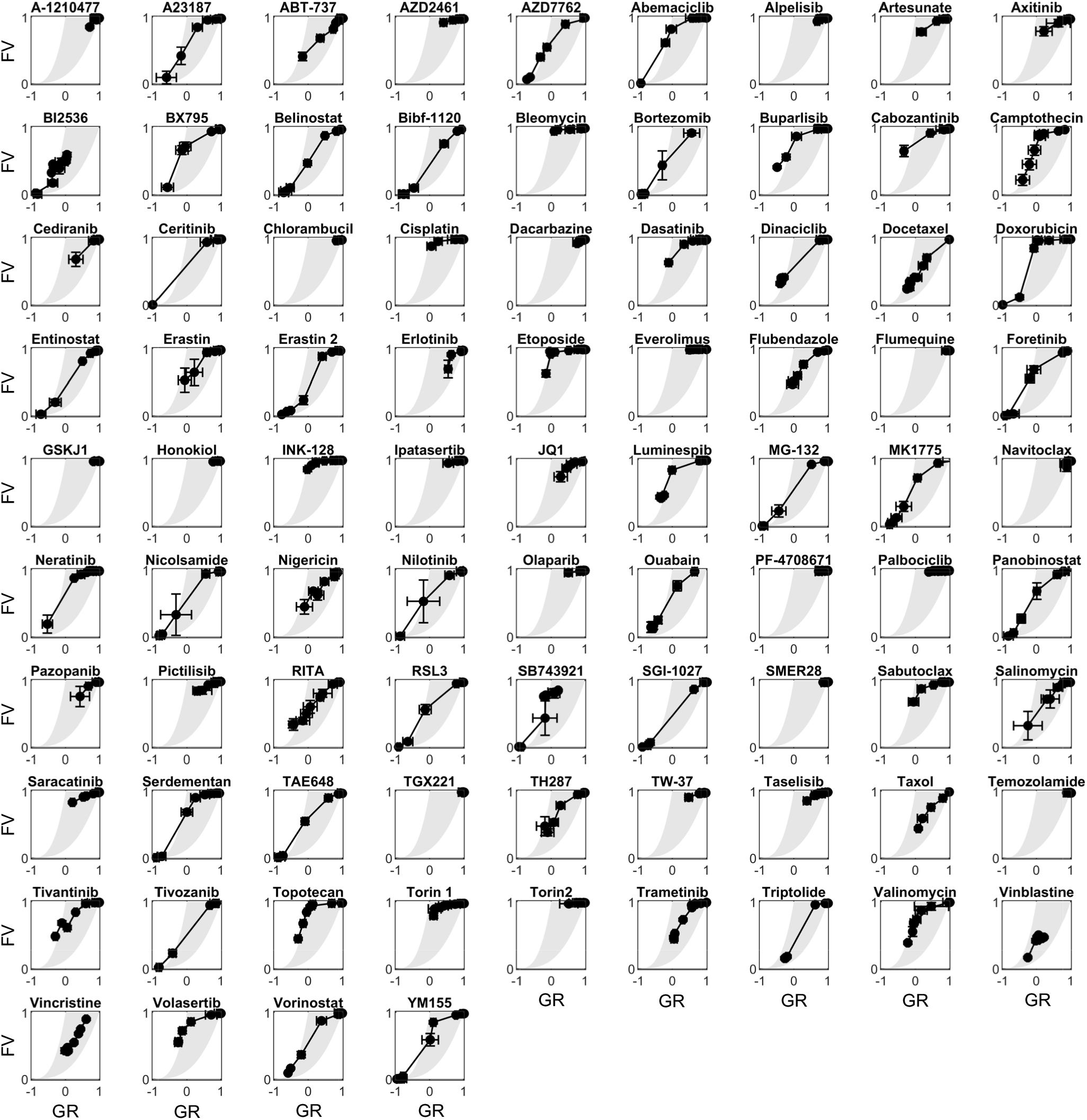
GR versus FV plots for all 85 drugs tested in U2OS. GR versus FV plots for 85 chemotherapies tested in U2OS. Error bars represent the standard deviation across 4 biological replicates.

**Figure S7:**
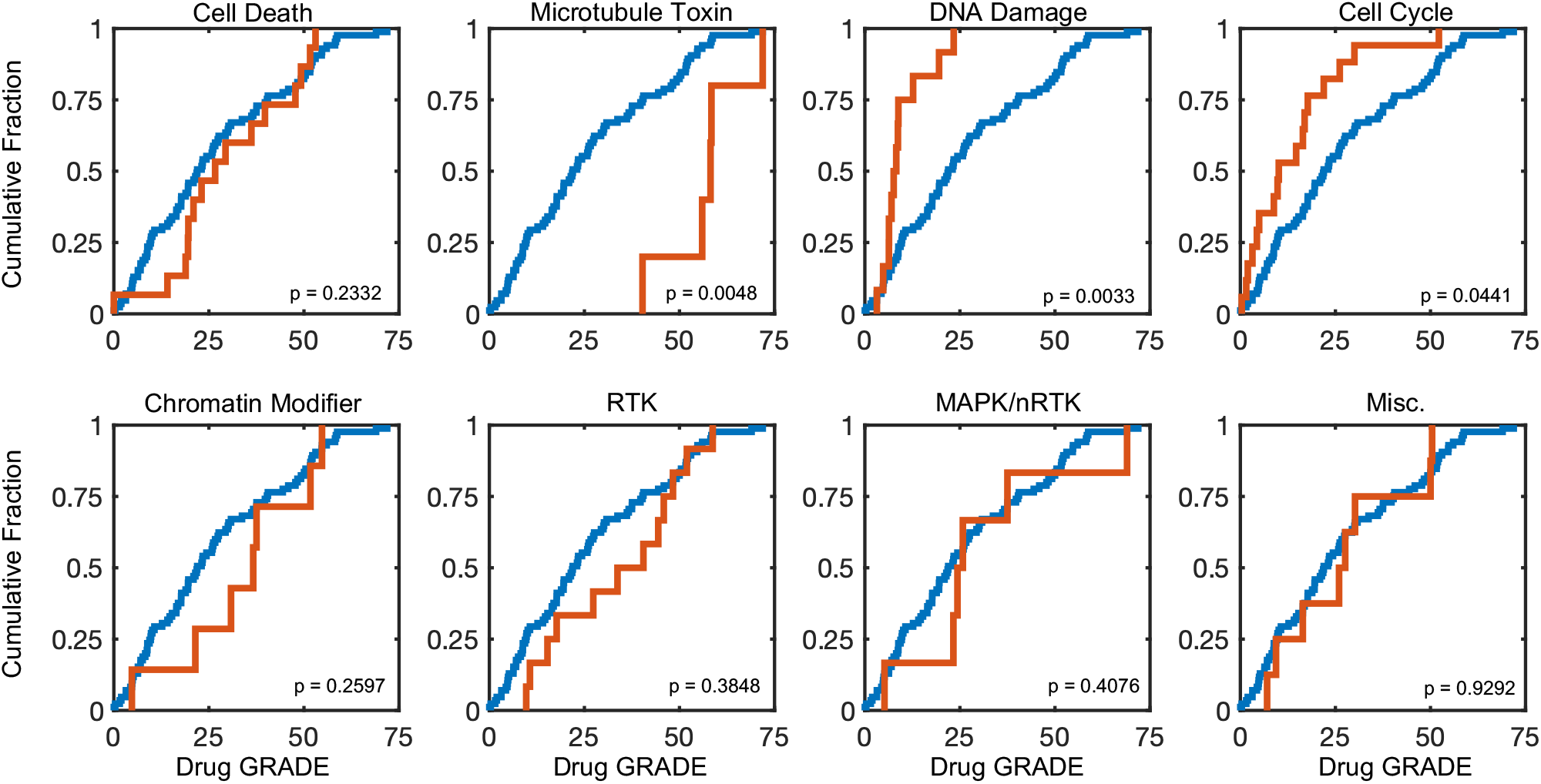
Drug GRADE varies by drug class. Cumulative distribution functions of Drug GRADE for all 85 drugs (blue) or drugs in the listed class (orange). Data are from U2OS cells. P values calculated using the a two-tailed KS test as in Figure 3E.

**Figure S8:**
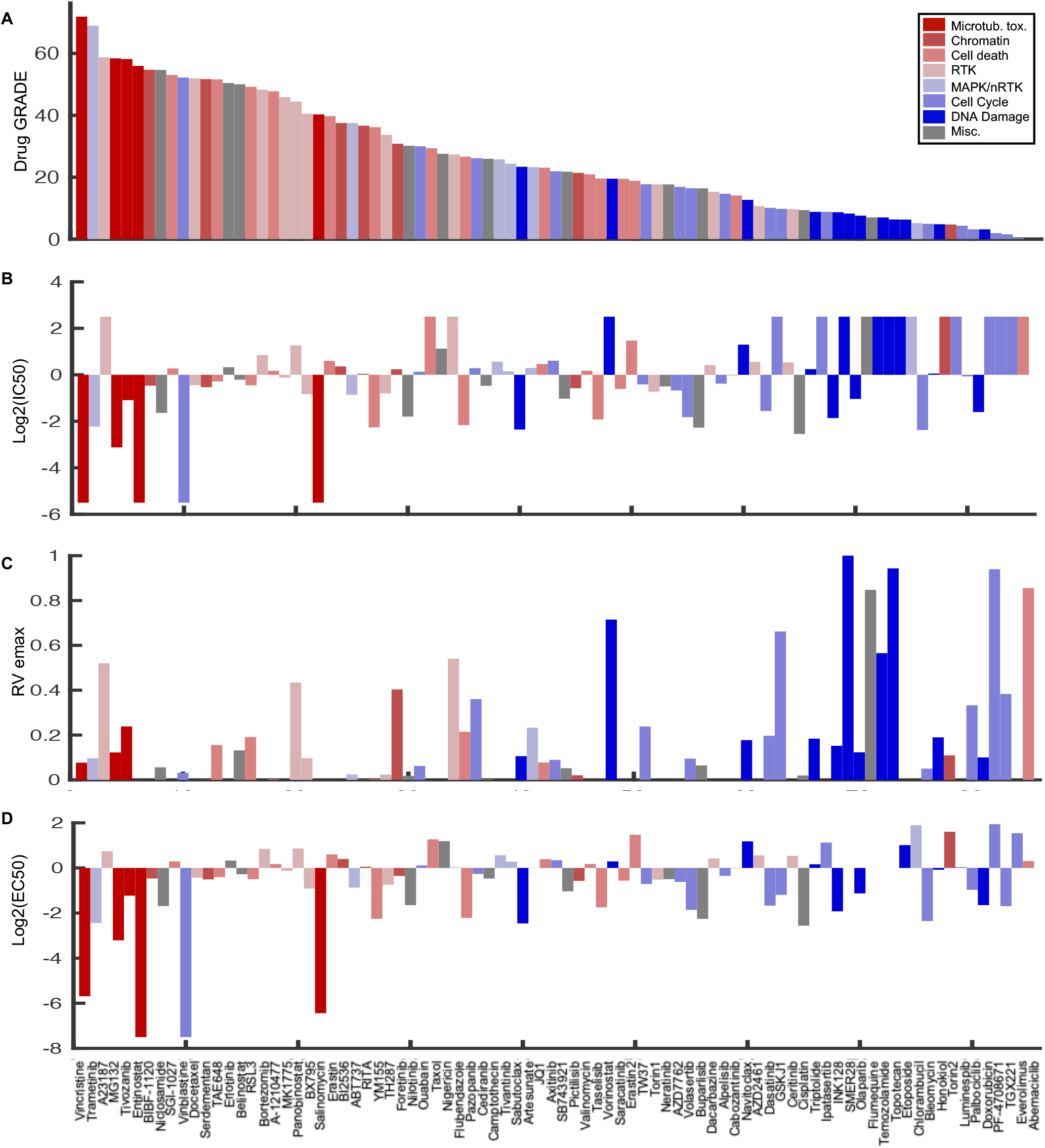
Relationship between Drug GRADE and traditional pharmaco-metrics. **(A)** Drug Grade shown for all 85 drugs profiled in this study. Data are colored by drug class as in Figure 3D. **(B)** Traditional IC50 (IC50 of the RV curve, i.e. “RV50”) **(C)** RV emax and **(D)** RV EC50. Data in (B-D) are in the same order as in (A) and colored as in (A).

**Figure S9:**
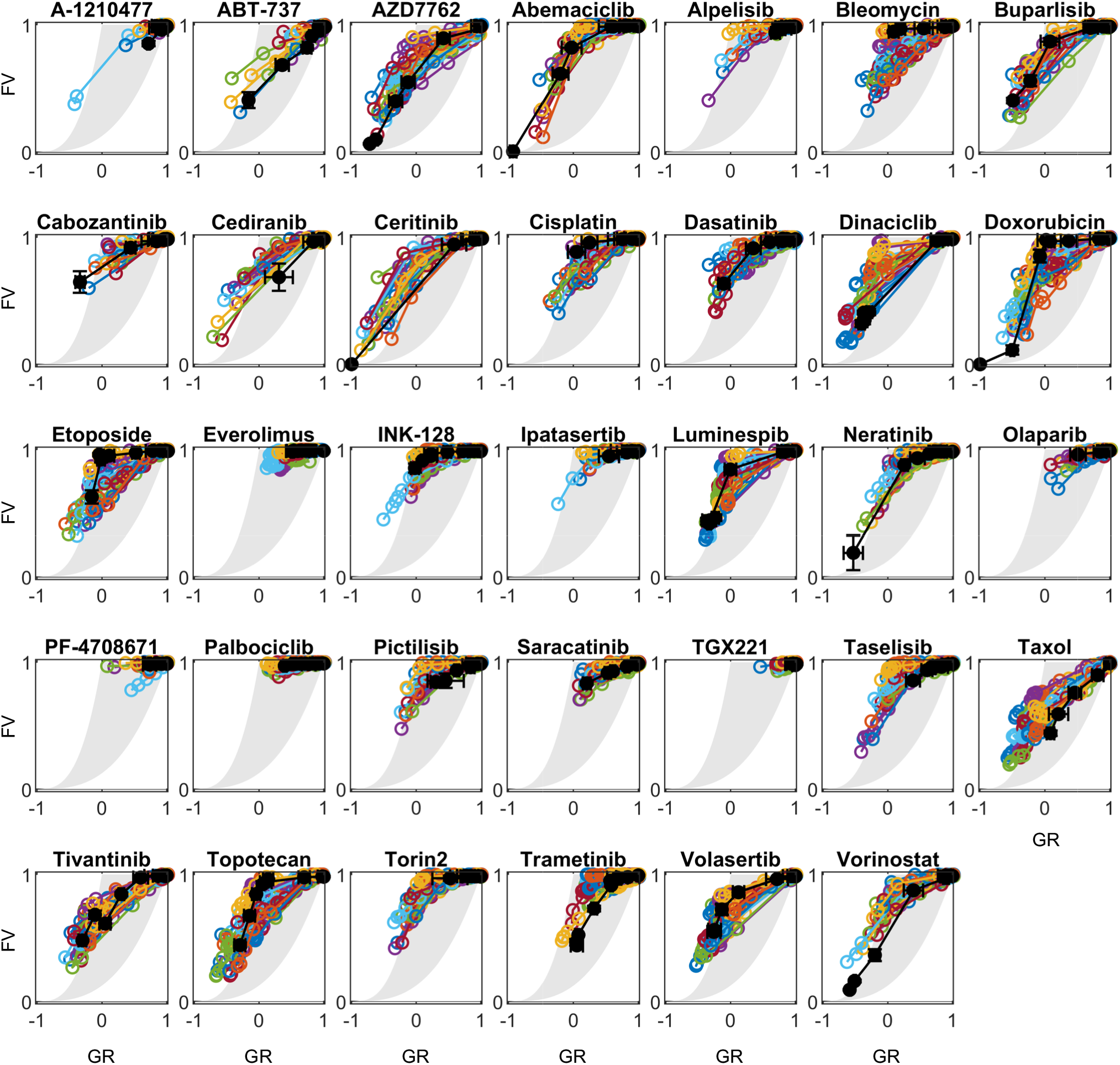
GR/FV for all drugs across U2OS and 35 LINCS cell lines. GR/FV plots for all 34 drugs tested by Hafner et al. across 24 LINCS consortium cell lines and U2OS. All LINCS consortium cell lines are shown as a colored dot and line. U2OS is shown as a black dot and line. Error bars for U2OS are the standard deviation across 4 replicates.

**Figure S10:**
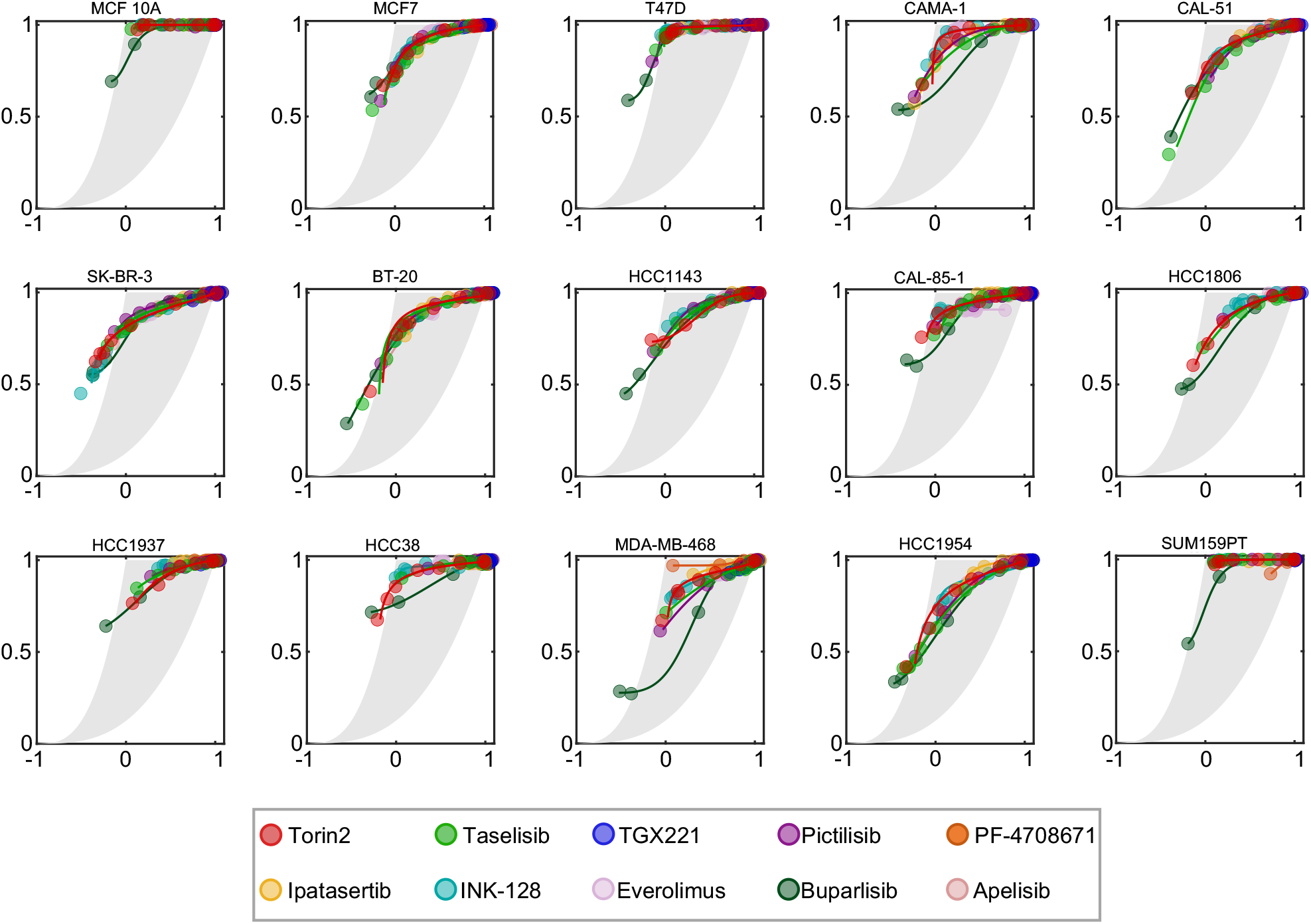
GR versus FV plots for all PI3K/mTOR targeting drugs from the LINCS consortium. GR/FV plots for 10 different chemotherapies that inhibit PI3K/mTOR in 15 cell lines from the LINC consortium.

## REFERENCES

Albeck, J.G., Burke, J.M., Aldridge, B.B., Zhang, M., Lauffenburger, D.A., and Sorger, P.K. (2008). Quantitative Analysis of Pathways Controlling Extrinsic Apoptosis in Single Cells. Mol Cell 30, 11–25.

Bruno, P.M., Liu, Y., Park, G.Y., Murai, J., Koch, C.E., Eisen, T.J., Pritchard, J.R., Pommier, Y., Lippard, S.J., and Hemann, M.T. (2017). A subset of platinum-containing chemotherapeutic agents kills cells by inducing ribosome biogenesis stress. Nat Med 23, 461–471.

Carey, L.A., Dees, E.C., Sawyer, L., Gatti, L., Moore, D.T., Collichio, F., Ollila, D.W., Sartor, C.I., Graham, M.L., and Perou, C.M. (2007). The Triple Negative Paradox: Primary Tumor Chemosensitivity of Breast Cancer Subtypes. Clin Cancer Res 13, 2329–2334.

Chopra, S.S., Jenney, A., Palmer, A., Niepel, M., Chung, M., Mills, C., Sivakumaren, S.C., Liu, Q., Chen, J.-Y., Yapp, C., et al. (2019). Torin2 Exploits Replication and Checkpoint Vulnerabilities to Cause Death of PI3K-Activated Triple-Negative Breast Cancer Cells. Cell Syst.

Fallahi-Sichani, M., Honarnejad, S., Heiser, L.M., Gray, J.W., and Sorger, P.K. (2013). Metrics other than potency reveal systematic variation in responses to cancer drugs. Nat Chem Biol 9, 708–714.

Forcina, G.C., Conlon, M., Wells, A., Cao, J.Y., and Dixon, S.J. (2017). Systematic Quantification of Population Cell Death Kinetics in Mammalian Cells. Cell Syst 4, 1–18.

Hafner, M., Niepel, M., Chung, M., and Sorger, P.K. (2016). Growth rate inhibition metrics correct for confounders in measuring sensitivity to cancer drugs. Nat Methods 13, 1–11.

Hafner, M., Mills, C.E., Subramanian, K., Chen, C., Chung, M., Boswell, S.A., Everley, R.A., Liu, C., Walmsley, C.S., Juric, D., et al. (2019). Multiomics Profiling Establishes the Polypharmacology of FDA-Approved CDK4/6 Inhibitors and the Potential for Differential Clinical Activity. Cell Chem Biol 26, 1067–1080.e8.

Haibe-Kains, B., El-Hachem, N., Birkbak, N.J., Jin, A.C., Beck, A.H., Aerts, H.J.W.L., and Quackenbush, J. (2013). Inconsistency in large pharmacogenomic studies. Nature 504, 389–393.

Harris, L.A., Frick, P.L., Garbett, S.P., Hardeman, K.N., Paudel, B.B., Lopez, C.F., Quaranta, V., and Tyson, D.R. (2016). An unbiased metric of antiproliferative drug effect in vitro. Nat Methods 13, 497–500.

Lachmann, A., Giorgi, F.M., Alvarez, M.J., and Califano, A. (2016). Detection and removal of spatial bias in multiwell assays. Bioinform Oxf Engl 32, 1959–1965.

Lin, A., Giuliano, C.J., Palladino, A., John, K.M., Abramowicz, C., Yuan, M.L., Sausville, E.L., Lukow, D.A., Liu, L., Chait, A.R., et al. (2019). Off-target toxicity is a common mechanism of action of cancer drugs undergoing clinical trials. Sci Transl Med 11, eaaw8412.

Méry, B., Guy, J.-B., Vallard, A., Espenel, S., Ardail, D., Rodriguez-Lafrasse, C., Rancoule, C., and Magné, N. (2017). In Vitro Cell Death Determination for Drug Discovery: A Landscape Review of Real Issues. J Cell Death 10, 1179670717691251.

Meyer, C.T., Wooten, D.J., Paudel, B.B., Bauer, J., Hardeman, K.N., Westover, D., Lovly, C.M., Harris, L.A., Tyson, D.R., and Quaranta, V. (2019). Quantifying Drug Combination Synergy along Potency and Efficacy Axes. Cell Syst 8, 1–44.

Overholtzer, M., Mailleux, A.A., Mouneimne, G., Normand, G., Schnitt, S.J., King, R.W., Cibas, E.S., and Brugge, J.S. (2007). A Nonapoptotic Cell Death Process, Entosis, that Occurs by Cell- in-Cell Invasion. Cell 131, 966–979.

Richards, R., Schwartz, H.R., Stewart, M.S., Cruz-Gordillo, P., Honeywell, M.E., Joyce, A.J., Landry, B.D., and Lee, M.J. (2019). Drug Combination Antagonism and Single Agent Dominance Result from Differences in Death Activation Kinetics. Biorxiv 805093.

Riss, T., Niles, A., Moravec, R., Karassina, N., and Vidugiriene, J. (2019). Cytotoxicity Assays_: n vitro methods to measure dead cells. p.

